# Hand–foot coordination is significantly influenced by motion direction in individuals with autism spectrum disorder

**DOI:** 10.1101/2022.03.30.486494

**Authors:** Yumi Umesawa, Kanae Matsushima, Reiko Fukatsu, Yasuo Terao, Masakazu Ide

## Abstract

Generally, when individuals attempt to move two limbs rhythmically in the opposite direction (e.g., flex the left hand and extend the left foot along the sagittal plane), the movements tend to be instead performed in the same direction. This phenomenon, known as directional constraint, can be harnessed to examine the difficulties in movement coordination exhibited by most individuals with autism spectrum disorder (ASD). While such difficulties have already been investigated through standardized clinical assessments, they have not been examined through kinematic methods. Thus, we employed a clinical assessment scale in an experimentally controlled environment to investigate whether stronger directional constraint during the rhythmic movement of two limbs is more pronounced and associated with decreased movement coordination in individuals with ASD. ASD and typically developing (TD) participants were asked to rhythmically move two limbs either in the same or opposite directions. In addition, the coordination skills of participants were assessed using the Bruininks-Oseretsky Test of Motor Proficiency Second Edition (BOT-2). Subjects with ASD showed significantly stronger directional constraint than TD participants during the contralateral and ipsilateral movement of the hand and foot. According to the pooled data from both groups, participants who showed stronger directional constraint during these two movement conditions also exhibited poorer coordinated movement skills in the BOT-2. Furthermore, stronger directional constraint was associated with severe autistic traits. These results suggest that people with ASD may have difficulties in inhibiting the neural signals that synchronize the direction of inter-limb movements, thus resulting in coordination disabilities.

**Lay Summary:** Individuals with autism spectrum disorder (ASD) often exhibit difficulties in coordinated movements. We asked those with ASD and healthy participants to move two limbs (e.g., left hand and left foot) either in the same or the opposite direction. Results demonstrated that participants with ASD had more difficulties in counteracting the tendency of their hand and foot to synchronously move in the same direction. Our findings suggested that difficulties to suppress synchronized movements of the hand and foot result in coordination disabilities.

## Introduction

Autism spectrum disorder (ASD) is an inclusive term for a group of neurodevelopmental disorders that share similar impairments in communication and reciprocal social interaction, with a tendency toward restricted, repetitive behavior (APA, 2013). Although the diagnostic criteria of autism focus on social impairments, many studies have reported that ASD is often comorbid with various motor disabilities, such as clumsiness, motor coordination abnormalities, postural control impairments, and poor performance in standardized tests of motor functioning (Fournier, Hass, Naik, Lodha, & Cauraugh, 2010; Green et al., 2009; Lim, Partridge, Girdler, & Morris, 2017; Molloy, Dietrich, & Bhattacharya, 2003; Paquet, Olliac, Bouvard, Golse, & Vaivre-Douret, 2016). Fournier et al. (2010) conducted a meta-analysis of 83 studies that focused on motor coordination in autism, and the results suggested substantial deficits in ASD across a wide range of behaviors. In addition, a demographic study reported that 79% of children with ASD had definite impairments in coordinated movement, as measured by the Movement Assessment Battery for Children (M-ABC), and 10% had borderline personality problems (Green et al., 2009). Previous studies have reported that such poorer coordination skills might cause difficulties in daily life (see the review by Gowen & Hamilton, 2013).

Previous studies have investigated the etiology of motor disabilities and have specifically focused on visuomotor coordination (Gowen & Hamilton, 2013; Sacrey, Germani, Bryson, & Zwaigenbaum, 2014). For instance, Haswell et al. (2009) found that children with ASD tend to rely more on proprioceptive feedback than visual feedback when learning reaching movements. On the contrary, typically developing (TD) individuals exhibited greater reliance on visual feedback (Haswell, Izawa, Dowell, Mostofsky, & Shadmehr, 2009). Additionally, our previous study revealed that individuals with ASD tended to show decreased utilization of visual landmarks to recognize the reaching position, while healthy individuals were more likely to use landmarks (Umesawa, Atsumi, Fukatsu, & Ide, 2020). Despite many reports of abnormal characteristics in individuals with ASD regarding visuomotor coordination (i.e., reaching movement), few studies have focused on the neural circuit underlying difficulties in inter-limb coordination.

Motor tasks in which TD individuals also exhibit difficulties in controlling their limbs as intended would provide valuable insights regarding the mechanisms underlying decreased coordination in ASD. One such task is the rhythmical movement of limbs in opposite directions (e.g., left hand flexion and left foot extension along the sagittal plane), during which these movements tend to be instead performed in the same direction, even in TD individuals (Baldissera, Cavallari, & Civaschi, 1982). This phenomenon is known as directional constraint and occurs because the neural excitability deriving from the movement of one limb modulates the other limb’s movement, with the overall intent to synchronize the direction of the two actions (Baldissera, Borroni, Cavallari, & Cerri, 2002). In addition, the degree of directional constraint differs depending on the combination of the limbs performing the action (Kelso & Jeka, 1992; Meesen, Wenderoth, Temprado, Summers, & Swinnen, 2006; Nakagawa, Muraoka, & Kanosue, 2015; Swinnen et al., 1997). In fact, while directional constraint is more evident during the concomitant movement of upper and lower limbs on the ipsilateral side (e.g., right hand and right foot), it is less prominent when moving an upper limb and a lower limb on the contralateral side (e.g., right hand and left foot). On the contrary, for coordinated bilateral movements of both limbs (e.g., both hands or both feet) along the sagittal plane, little or no directional constraint is observed (Carson & Riek, 2001; Nakagawa et al., 2015).

Several studies have investigated the brain areas responsible for directional constraint. For example, Byblow et al. (2007) examined the functional connectivity between the secondary and primary motor areas during active ankle dorsiflexion and plantarflexion using dual-coil paired-pulse transcranial magnetic stimulation (TMS) (Byblow et al., 2007). The results indicated that neural networks involving the premotor area (PMA) and primary motor area (M1) facilitate hand–foot movements in the same direction. Steyvers et al. (2003) used repeated TMS to suppress the supplementary motor area (SMA) and evaluated the performance of in-phase and anti-phase bimanual index finger movements. After stimulation, the mean relative phase error between the hands only increased in the anti-phase trials (Steyvers et al., 2003). Additionally, anodal transcranial direct current stimulation (tDCS) of the SMA, which increases the excitability of this area, could improve the accuracy and stability of anti-phase bimanual supination-pronation movements (Carter, Maslovat, & Carlsen, 2015). Together, these findings suggest that the secondary motor areas (PMA and SMA) may play a key role in determining the degree of directional constraint during inter-limb coordinated movements. Additionally, these findings indicate that neural activation in the PMA facilitates synchronization of inter-limb movements, while those in the SMA contribute to the execution of anti-phase movements and inhibit directional constraint.

One representative feature of ASD is a severe excitability/inhibitory imbalance caused by alterations in gamma-aminobutyric acid (GABA) levels (Pizzarelli & Cherubini, 2011). GABA is a major cortical inhibitory neurotransmitter that depresses neural activity in the cerebral cortex (Krnjevic & Schwartz, 1967). In our previous work, we measured and compared GABA concentration in the M1 and SMA of subjects with ASD and healthy individuals using ^1^H-magnetic resonance spectroscopy (^1^H-MRS) (Umesawa et al., 2020), which is a non-invasive neuroimaging method that can estimate the concentrations of specific chemical metabolites and neurotransmitters in vivo (Jansen, Backes, Nicolay, & Kooi, 2006). We found that a lower GABA concentration in the SMA was associated with poorer coordinated movement skills in ASD participants. Based on these findings, it is possible that decreased GABA levels in the SMA cause strong directional constraints in individuals with ASD.

In this study, we investigated whether a stronger directional constraint is observed in individuals with ASD during the rhythmic movement of two limbs. We also elucidated whether decreased coordination skills as evaluated through a standardized assessment tool are associated with a higher degree of directional constraint. Specifically, our study focused on phase transitions during cyclic inter-limb movements and aimed to clarify kinematic features in an experimentally controlled situation. The findings of this study will shed new light on the complex mechanisms of motor difficulties in autism.

## Methods

### Participants

The present study was conducted according to the guidelines of the Declaration of Helsinki and approved by the Ethics Committee of the National Rehabilitation Center for Persons with Disabilities. All participants and their parents provided written informed consent after the study procedures were fully explained.

We recruited 25 individuals with ASD and 23 age-matched TD individuals. Individuals with clinically diagnosed ASD were recruited through support groups aimed at parents of children with developmental disorders and from the Department of Child Psychiatry at the Hospital of National Rehabilitation Center for Persons with Disabilities (Saitama, Japan). Participants completed the Japanese version of the Autism Spectrum Quotient (AQ) scale (Baron-Cohen, Wheelwright, Skinner, Martin, & Clubley, 2001; Wakabayashi, Tojo, Baron-Cohen, & Wheelwright, 2004), in which higher scores indicate more severe autistic traits. Intelligence quotients (IQs) were assessed using the Wechsler Adult Intelligence Scale-Third Edition (WAIS-III). The Wechsler Intelligence Scale for Children-Fourth Edition (WISC-IV) was also used to evaluate two ASD participants below the age of 15 years. Handedness was assessed through the laterality quotient score of the Edinburgh Handedness Inventory (Oldfield, 1971).

### Procedure

For 11 of the participants with ASD and five of the TD participants, hand–foot coordination and motor skills were assessed on the same day, while for the remainder of the subjects (14 participants with ASD and 18 TD participants), the two assessments were completed on different days.

#### (a) Measurement of hand–foot coordination movements

The participants sat comfortably in a chair with the right and left forearm fixed in a prone position on an armrest. The knee was supported using the soft pad with which the chair was equipped. The knee and elbow were positioned at 130° and 90° of flexion, respectively. During the execution of the tasks, the subjects kept their eyes closed and wore headphones. Kinematic data were collected using a 5-camera, 3D motion capture system (OptiTrack, Acuity Inc., Japan) at a frequency of 100 Hz (spatial resolution was less than 0.5 mm). The system registered the 3D trajectories of the passive markers. The markers were attached to the fifth metacarpal and metatarsal heads.

During the same (SAME) and opposite (OPP) directional movement tasks, the two limbs were moved along the sagittal plane in the same or opposite directions, respectively. Three limb combinations were assessed: the right and left hand, right hand and left foot, and right hand and right foot (Fig. 1). In order to avoid laterality-related effects, participants were asked to perform the movements with the opposite hand and foot in half of the trials in the bilateral and ipsilateral conditions (i.e., left hand and right foot, left hand and left foot). Subjects moved the two limbs for approximately 30 s per trial at the pace set by the constant rhythm (1.5 Hz) of a metronome. Each task consisted of six trials, resulting in a total of 36 trials (two directions × three combinations × six trials). The order of conditions was randomized. Each task took approximately 60 min to complete, including the practice trials.

**Figure 1.**
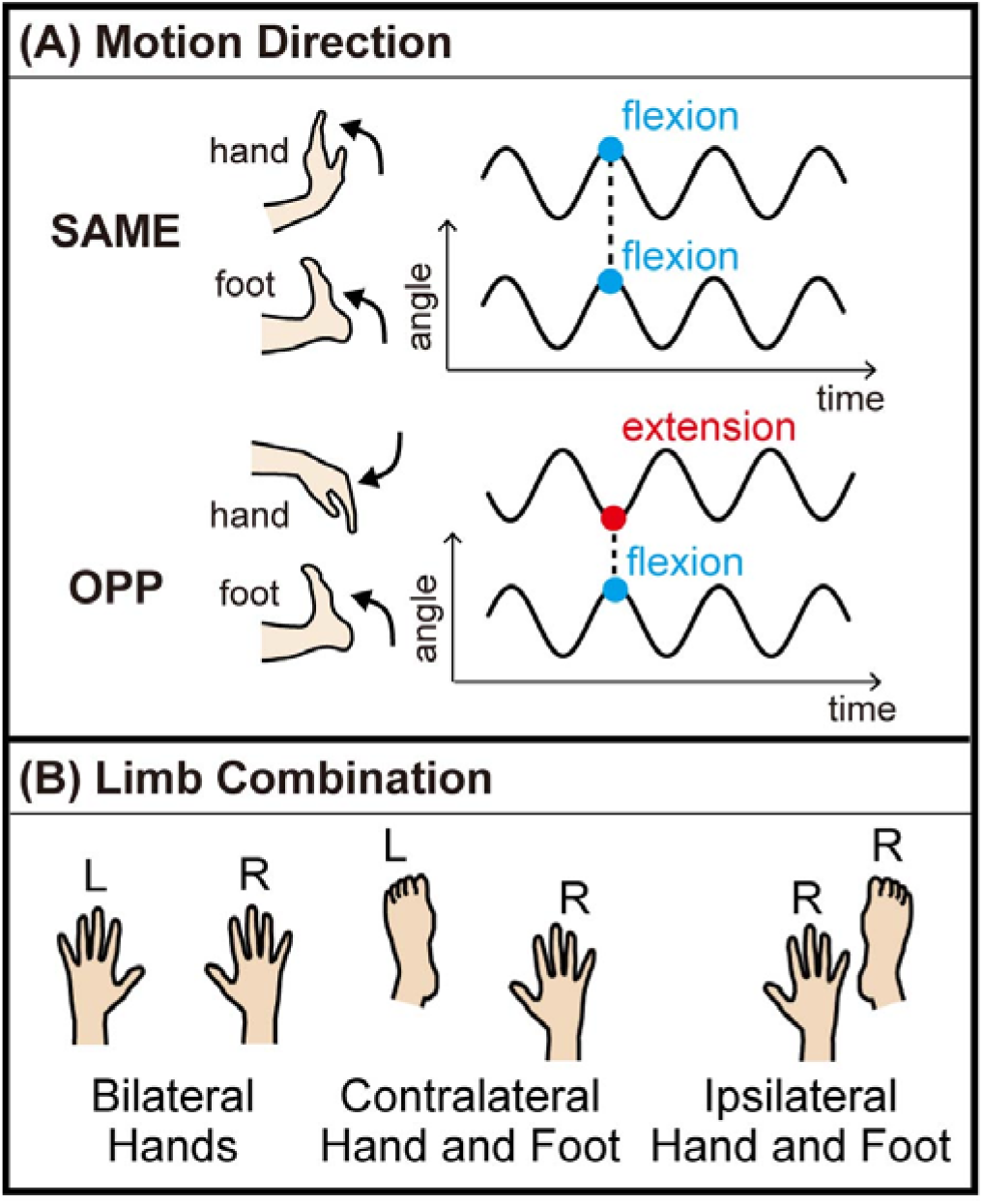
Summary of experimental conditions. (A) The two movement direction conditions included movements in the same (SAME) and opposite (OPP) direction. (B) The three limb combination conditions included the bilateral movement of the hands (left), the contralateral movement of the hand and foot (center), and the ipsilateral movement of the hand and foot (right).

#### (b) Assessment of motor skills

We used the Bruininks-Oseretsky Test of Motor Proficiency second edition (BOT-2) to evaluate the motor skills of each participant. The BOT-2 is the most widely used standardized measure of the motor skills that are impaired in developmental coordination disorders. This test examines both fine and gross motor skills (Crowe, 1989; Robert & Brett, 2005). With regard to the former, the BOT-2 assesses precise bodily control requiring finger and hand movements based on visuomotor integration (fine manual control) and bimanual/arm-hand coordination (manual coordination). The gross motor skills assessed by the BOT-2 include maintaining posture, sequential and simultaneous body coordination (body coordination), and strength of the trunk and the upper and lower body (strength and agility). For each category (i.e., fine manual control, manual coordination, body coordination, strength, and agility), scores were measured according to the participant’s performance. In addition, each of these categories is subdivided into two subsets, which in turn include several items, forming a total of eight subsets and 53 items. Specifically, the fine manual control category consists of fine motor precision (seven items) and fine motor integration (eight items); manual coordination includes manual dexterity (five items) and upper limb coordination (seven items); body coordination comprises bilateral coordination (seven items) and balance (nine items); and strength and agility includes running speed and agility (five items) and strength (five items).

### Kinematic data analysis

To evaluate the performance of two-limb coordination, the relative phase (Φ) between the movements of the two limbs was calculated for each cycle as follows:

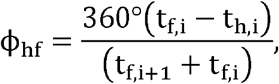

where t_h,i_ and t_f,i_ indicate the time of the ith peak extension of the hand and foot, respectively (Carson, Goodman, Kelso, & Elliott, 1995; Ridderikhoff, Peper, & Beek, 2005; Volman, Laroy, & Jongmans, 2006). Python 3.0 (Python Software Foundation, Wilmington, DE, USA) with the SciPy package (https://scipy.org) (Virtanen et al., 2020) was used to detect the time at which peak extension of the hand and foot was reached. To evaluate the accuracy of the coordinated movements, the absolute errors of the relative phases (AEФ) between the two limbs were computed. The AEΦ was calculated as the absolute value of the averaged errors in one trial to the target relative phase (SAME: 0°, OPP: 180°). We defined the degree of directional constraint through two different indexes obtained by calculating the difference between the performances of SAME and OPP conditions ([AEФ of OPP -AEФ of SAME]), as described in a previous study (Nakagawa et al., 2015). Each index was calculated using the data obtained from the 1st to the 20th cycle according to the following formula:

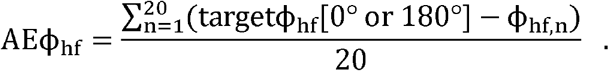

### Statistical analysis

The raw BOT-2 scores were standardized using the following formula:

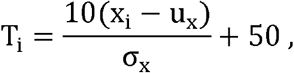

where x_i_ represents the sampled scores, and u_x_ and s_x_ correspond to the arithmetic mean and standard deviation of the scores for each of the four test categories, respectively. A two-way repeated measures analysis of variance (2 × 2 ANOVA) was conducted to compare the two groups and limb movement combinations (hands, bilateral, and ipsilateral). The Shaffer’s Modified Sequentially Rejective Bonferroni (MSRB) procedure was used for multiple comparisons. The Cohen’s d and partial η^2^ were calculated to assess the effect sizes of the intergroup difference and two-way ANOVA, respectively. We used SPSS Version 23.0 (IBM, Armonk, NY, USA) to perform the t-test and ANOVA and G*power 3.1 (https://www.psychologie.hhu.de/arbeitsgruppen/allgemeine-psychologie-und-arbeitspsychologie/gpower) to calculate the effect sizes.

## Results

### Demographic and clinical characteristics

The mean age was not different between the ASD and TD groups (two-tailed t-test: t (46) = 0.45, p = 0.66, Cohen’s d = 0.13; Table 1). AQ scores were significantly higher in the ASD group than in the TD group (t (46) = 6.72, p < 0.001, Cohen’s d = 1.94). The full-scale IQ (FIQ) and performance IQ (PIQ) were significantly different between the two groups (FIQ: t (46) = -2.76, p = 0.009, Cohen’s d = 1.07; PIQ: t (46) = -2.54, p = 0.015, Cohen’s d = 1.08). There were no significant between-group differences in age or verbal IQ (VIQ) (t (46) = -1.22, p = 0.228, Cohen’s d = 0.36) and in handedness (t (46) = -0.98, p = 0.330, Cohen’s d = 0.29).

**Table 1.** Participants’ information

|  | ASD group | TD group |
| --- | --- | --- |
| Sex (M:F) | 13:12 (N = 25) | 10:13 (N = 23) |
| Age, mean years (range) | 23.9 ($\pm 5.4$) | 23.3 ($\pm 3.9$) |
| LQ, mean (range) | 61.8 ($\pm 53.2$) | 75.7 ($\pm 43.8$) |
| AQ, mean (range)** | 33.3 ($\pm 7.7$) | 18.1 ($\pm 8.0$) |
| VIQ, mean (range) | 112.6 ($\pm 16.5$) | 117.9 ($\pm 12.6$) |
| PIQ, mean (range)* | 104.2 ($\pm 15.8$) | 114.8 ($\pm 12.3$) |
| FIQ, mean (range)** | 107.0 ($\pm 16.8$) | 118.0 ($\pm 10.5$) |

Abbreviations: M, male; F, female; ASD, autism spectrum disorder; TD, typically developing; LQ, laterality quotient; AQ, autism spectrum quotient; VIQ, verbal intelligence quotient; PIQ, performance intelligence quotient; FIQ, full scale intelligence quotient. AQ scores were evaluated using the AQ scale. The LQ scores were assessed using the Edinburgh Handedness Inventory (Oldfield, 1971). IQ was assessed using the Wechsler Adult Intelligence-Third Edition (WAIS-III). \* $p < 0.05$ , \*\* $p < 0.01$ .

### Effect of directional constraint

#### (1) Analysis of the AEФ

Figure 2 illustrates the AEΦ of the SAME and OPP directional movements for each limb combination condition. According to the repeated measures ANOVA, a main effect of limb combination was observed, but there was no main effect of group or interactions for movements in the SAME direction (group: F (1, 46) =1.27, p = 0.265, generalized η^2^ = 0.017; combination: F (1, 46) = 86.2, p < 0.001, generalized η^2^ = 0.41; interaction: F (1, 46) =0.47, p = 0.630, generalized η^2^ = 0.0038; Fig. 1A). Multiple comparisons revealed significantly increased values of AEΦ in contralateral and ipsilateral combinations compared with bilateral hand combinations (contralateral vs. bilateral: t (46) = 10.3, p < 0.001, Cohen’s d = 1.50; ipsilateral vs. bilateral: t (46) = 9.5, p < 0.001, Cohen’s d = 1.39). By contrast, main effects of group and limb combination were found for movements in OPP directions (group: F (1, 46) =12.3, p = 0.001, generalized η^2^ = 0.13; combination: F (1, 46) = 70.6, p < 0.001, generalized η^2^ = 0.40; Fig.1B). Furthermore, there was a significant interaction between group and limb combination (F (1, 46) =7.3, p = 0.001, generalized η^2^ = 0.063). Simple main effects analysis demonstrated that the AEФ was significantly different between the groups for movements in OPP directions in the contralateral and ipsilateral hand–foot combinations (contralateral: F (1, 46) = 10.0, p = 0.003, generalized η^2^ = 0.18; ipsilateral: F (1, 46) = 10.4, p = 0.002, generalized η^2^ = 0.18), while the intragroup difference in AEФ in the bilateral hand combination was not significant (F (1, 46) = 3.4, p = 0.07, generalized η^2^ = 0.069). We also found a simple main effect of limb combination in both groups (ASD: F (2, 48) = 56.6, p < 0.001, generalized η^2^ = 0.48; TD: F (2, 44) = 18.5, p < 0.001, generalized η^2^ = 0.29).

**Figure 2.**
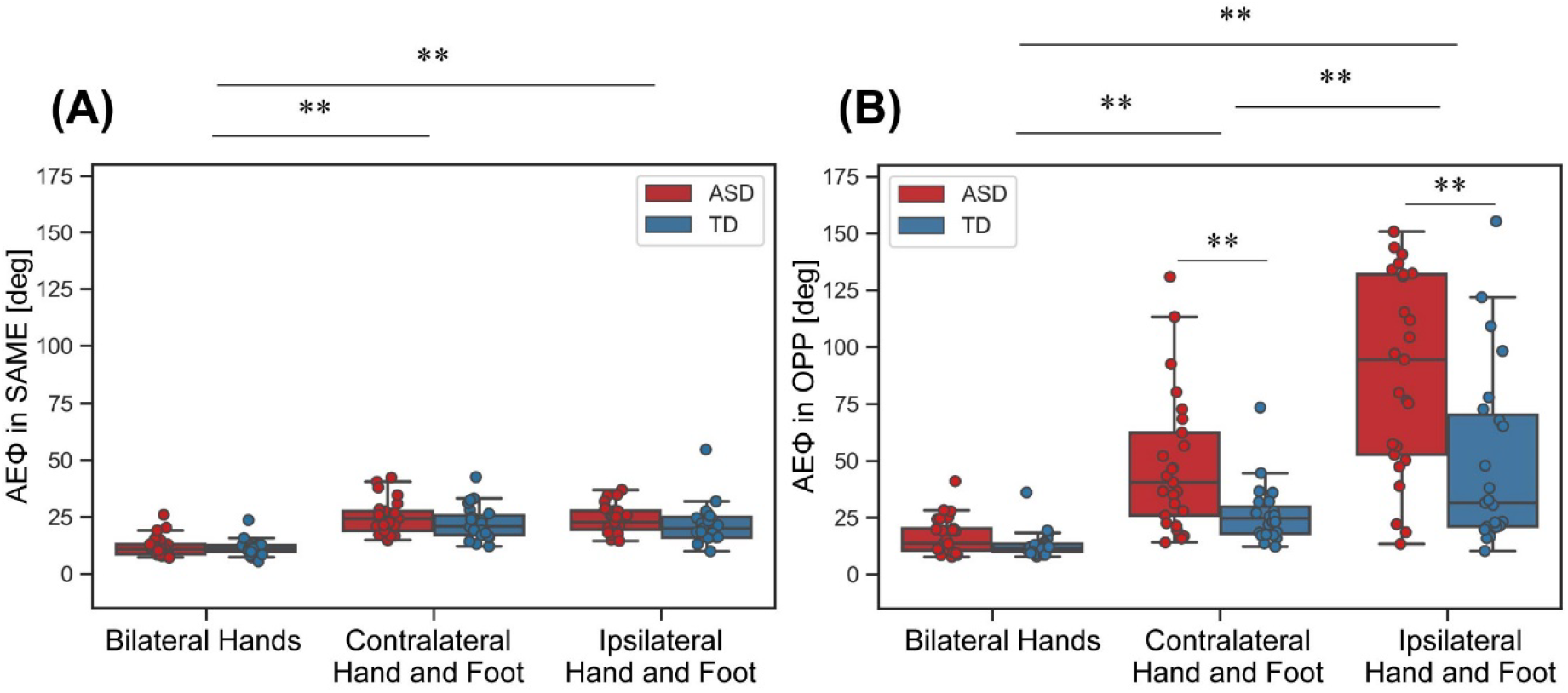
The absolute errors of the relative phases (AEФ) between the bilateral hand, contralateral hand and foot, and ipsilateral hand and foot conditions. (A) The AEФ of the ASD and TD groups for movements in the same direction (SAME). The upper and lower boundaries of the standard boxplots indicate the 25^th^ and 75^th^ percentiles. The horizontal line across the box marks the median of the distribution. The vertical lines below and above the box represent the minimum and maximum, respectively. (B) The AEФ of the ASD and TD groups for movements in opposite directions (OPP). **p < 0.01. Abbreviations: ASD, autism spectrum disorder; TD, typically developing.

#### (2) Analysis of the index of directional constraint

The degree of directional constraint was calculated by subtracting the AEФ in the SAME directional condition from the AEФ in the OPP directional condition, as previously described (Nakagawa et al., 2015), for each limb combination condition for the ASD and TD groups (Fig. 3). The two-way repeated measures ANOVA showed significant main effects and an interaction of group and limb combination (group: F (1, 46) = 13.0, p < 0.001, generalized η^2^ = 0.14; limb combinations: F (1, 46) = 57.3, p < 0.001, generalized η^2^ = 0.35; interaction: F (1, 46) = 8.0, p < 0.001, generalized η^2^ = 0.07). Multiple comparisons showed significant group differences in directional constraint among all limb combinations (bilateral – contralateral: t (46) = 3.9, p < 0.001, Cohen’s d = 0.65; bilateral – ipsilateral: t (46) = 8.2, p < 0.001, Cohen’s d = 1.45; contralateral – ipsilateral: t (46) = 7.7, p < 0.001, Cohen’s d = 0.97). Simple main effect analysis revealed that directional constraint in the ASD group was significantly greater than that in the TD group in the contralateral and ipsilateral combinations (contralateral: F (1, 46) = 10.6, p = 0.002, generalized η^2^ = 0.19; ipsilateral: F (1, 46) = 11.1, p = 0.002, generalized η^2^ = 0.19), while directional constraint in the bilateral combination was not significantly different between the groups (F (1, 46) = 3.0, p = 0.09, generalized η^2^ = 0.061). A simple main effect of limb combination was also found in both groups (ASD: F (2, 48) = 50.6, p < 0.001, generalized η^2^ = 0.46; TD: F (2, 44) = 12.9, p < 0.001, generalized η^2^ = 0.23). Notably, participants with ASD demonstrated a significantly higher degree of directional constraint when they performed cyclic movements in the contralateral hand–foot combination than in the bilateral hand combination, while the TD group showed no difference between the bilateral hand and contralateral hand–foot combination (ASD: t (24) = 4.0, p < 0.001, Cohen’s d = 0.98; TD: t (22) = 0.9, p = 0.36, Cohen’s d = 0.25).

**Figure 3.**
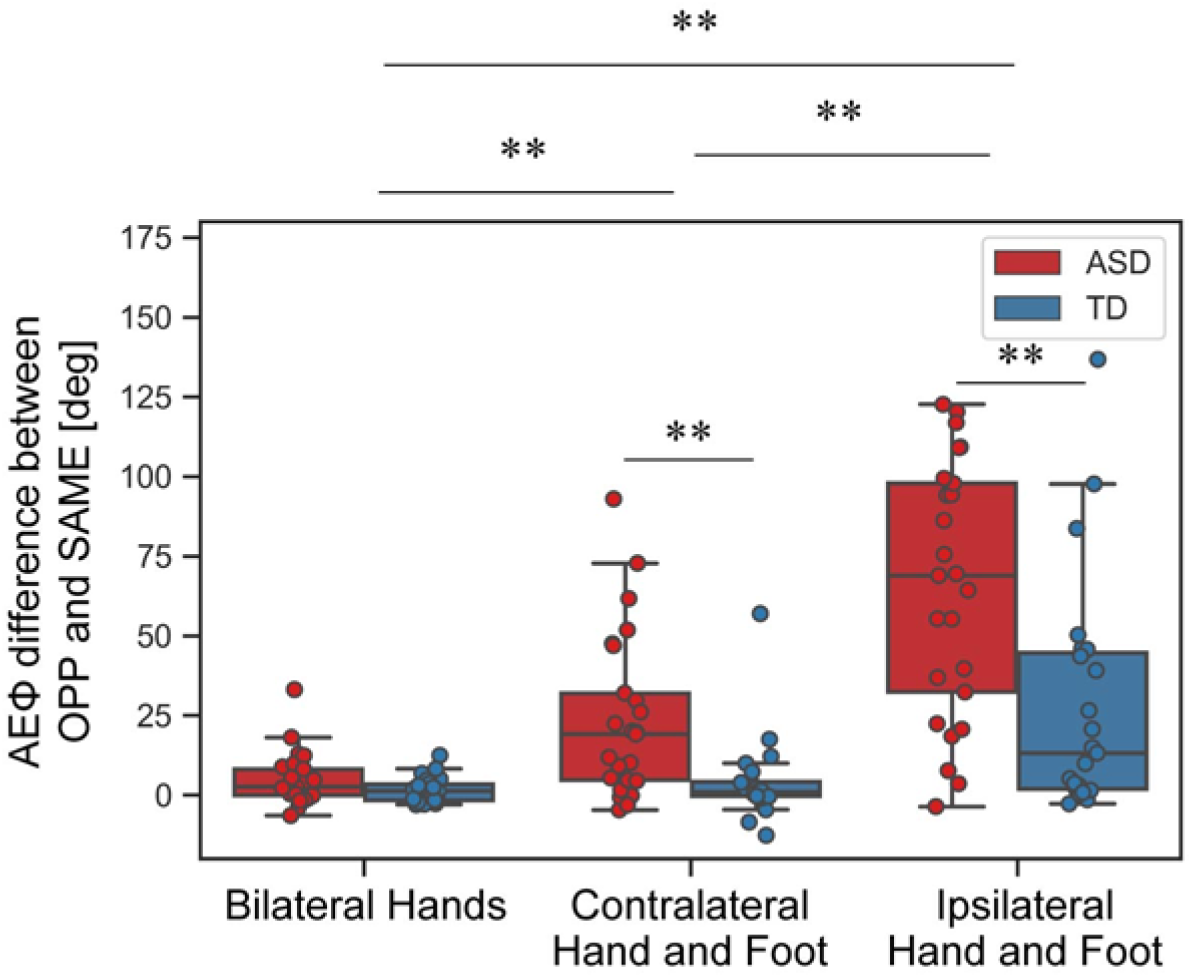
The degree of directional constraint between the bilateral hand, contralateral hand and foot, and ipsilateral hand and foot conditions. Directional constraint was calculated by subtracting the AEФ of the SAME from the AEФ of the OPP. **p < 0.01. Abbreviations: AEФ, absolute error of the relative phase; SAME, movement in the same direction; OPP, movement in opposite directions; ASD, autism spectrum disorder; TD, typically developing.

### Correlation between directional constraint and BOT-2 scores

The ASD group showed lower average scores in all four BOT-2 categories compared with the TD group (fine motor control: t (46) = -2.8, p = 0.004, Cohen’s d = 0.81; manual coordination: t (46) = -3.2, p = 0.001, Cohen’s d = 0.94; body coordination: t (46) = -3.9, p < 0.001, Cohen’s d = 1.16; strength and agility: t (46) = -3.8, p < 0.001, Cohen’s d = 1.1; Fig. 4). Figure 5 shows the relationship between the degree of directional constraint and the scores for each of the four BOT-2 categories. A Bonferroni correction for multiple comparisons of the BOT-2 scores was applied to adjust the alpha threshold to p < 0.0125.

**Figure 4.**
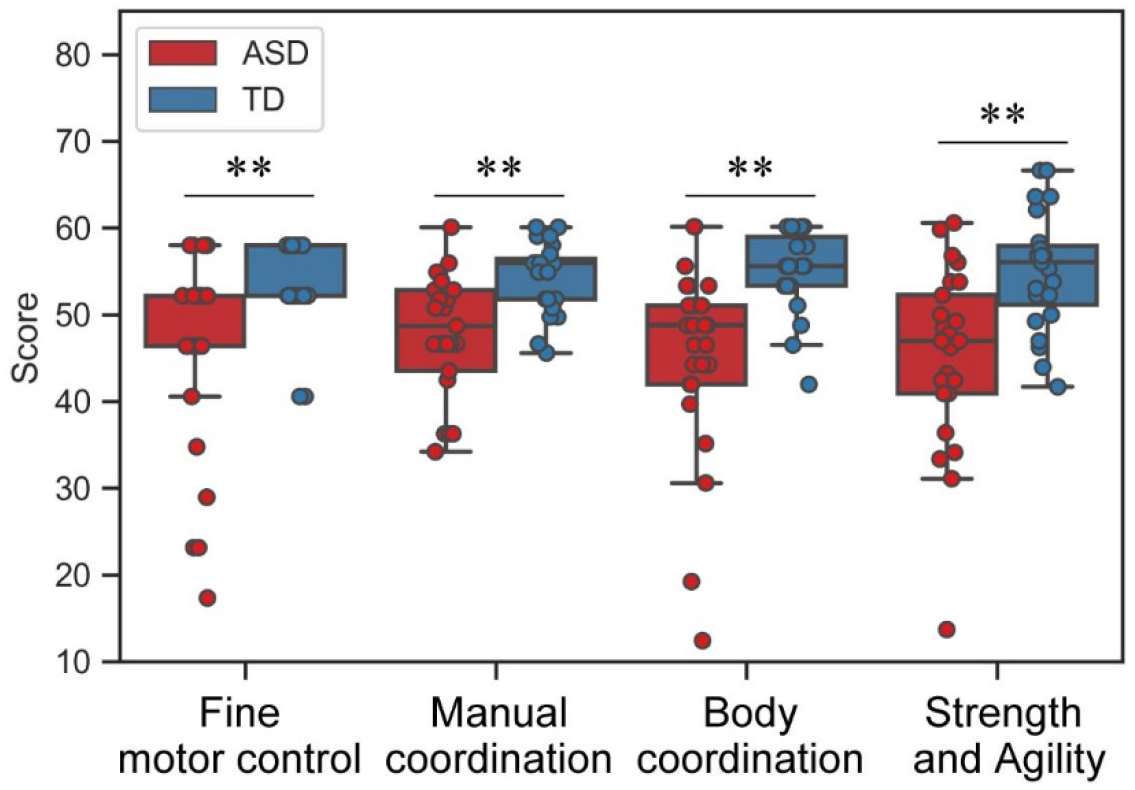
The BOT-2 scores for each of the four test categories in the ASD and TD groups. The upper and lower boundaries of the standard boxplots indicate the 25^th^ and 75^th^ percentiles. The horizontal line across the box marks the median of the distribution. The vertical lines below and above the box represent the minimum and maximum, respectively. **p < 0.01. Abbreviations: ASD, autism spectrum disorder; TD, typically developing; BOT-2, Bruininks-Oseretsky Test of Motor Proficiency second edition.

**Figure 5.**
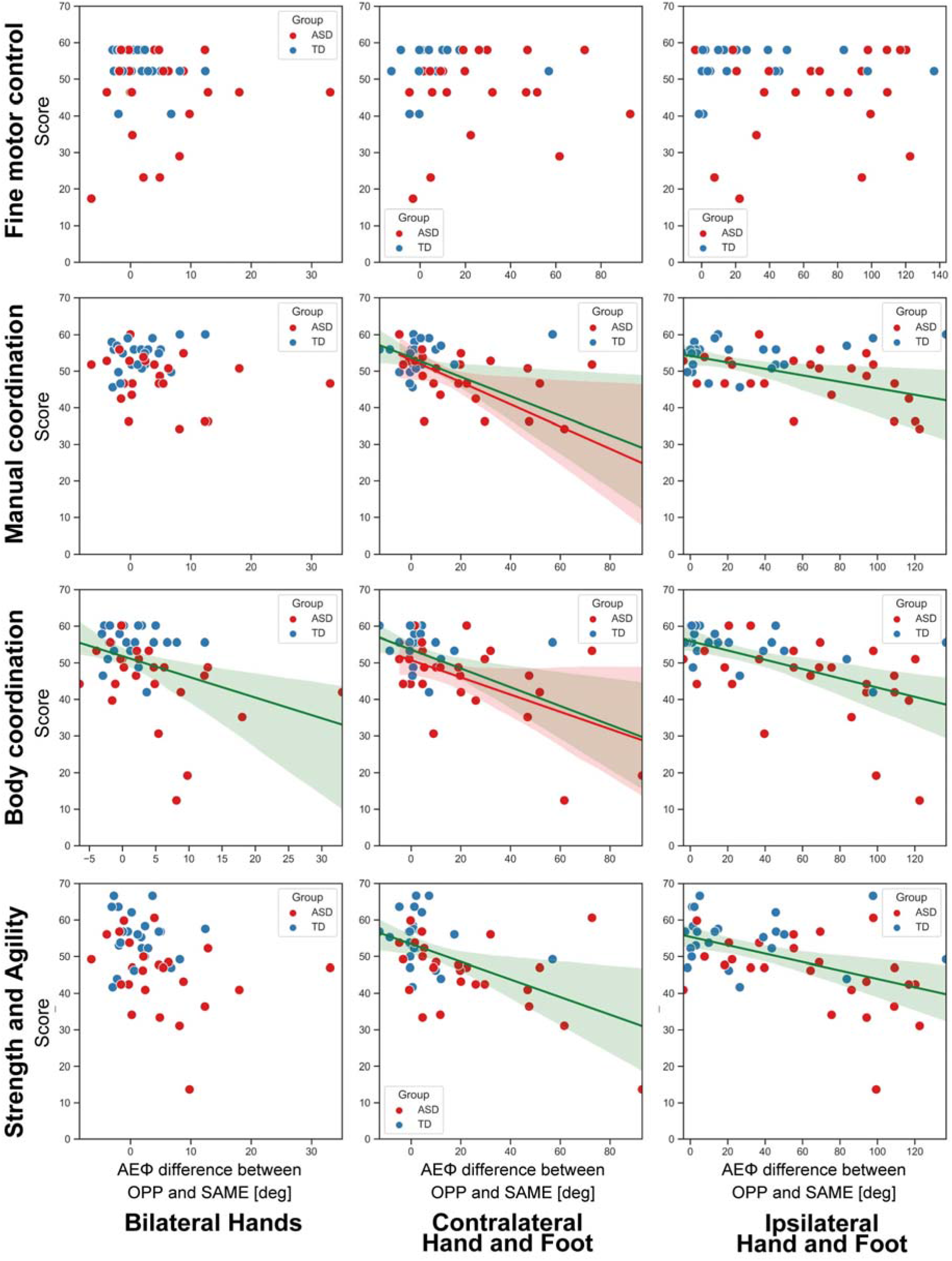
Correlation between the degree of directional constraint and BOT-2 scores. The shaded areas (green, red, and blue for ASD + TD, ASD, and TD, respectively) denote the 95% confidence intervals for the correlation. The horizontal axis indicates the degree of directional constraint in each of the three limb combinations (bilateral hand, contralateral hand and foot, and ipsilateral hand and foot). The vertical axis indicates the BOT-2 scores in each of the four test categories (fine manual control, manual coordination, body coordination and strength and agility). Abbreviations: ASD, autism spectrum disorder; TD, typically developing; BOT-2, Bruininks-Oseretsky Test of Motor Proficiency second edition; AEФ, absolute error of the relative phase; SAME, movement in the same direction; OPP, movement in opposite directions.

There was no significant correlation between the fine motor control scores and directional constraint for all limb combinations. Manual coordination scores were significantly correlated with directional constraint in the contralateral and ipsilateral combinations (contralateral: r = - 0.60, p < 0.001, power (1 – β) = 0.99; ipsilateral: r = - 0.37, p = 0.009, power (1 – β) = 0.75). Body coordination scores were significantly negatively correlated with the degree of directional constraint for all limb combinations (bilateral: r = - 0.38, p = 0.008, power (1 – β) = 0.77; contralateral: r = - 0.58, p < 0.001, power (1 – β) = 0.99; ipsilateral: r = - 0.54, p < 0.001, power (1 – β) = 0.98). Moreover, strength and agility scores were negatively correlated with the degree of directional constraint in the contralateral and ipsilateral combinations (contralateral: r = - 0.55, p < 0.001, power (1 – β) = 0.99; ipsilateral: r = - 0.48, p < 0.001, power (1 – β) = 0.94).

We also analyzed the correlation between BOT-2 scores and directional constraints for each group. We found that directional constraint was significantly correlated with manual coordination scores and body coordination scores, but only in the ASD group for the contralateral condition (manual coordination: r = - 0.64, p < 0.001, power (1 - β) = 0.95; body coordination: r = - 0.54, p = 0.005, power (1 - β) = 0.83).

### Correlation between directional constraint and AQ scores

We investigated the relationship between the degree of directional constraint and autistic traits (i.e., AQ score). Figure 6 shows that participants with higher AQ scores showed a greater degree of directional constraint in all limb combinations (bilateral: r = 0.34, p = 0.020, power (1 - β) = 0.67; contralateral: r = 0.48, p < 0.001, power (1 - β) = 0.94; ipsilateral: r = 0.44, p = 0.002, power (1 - β) = 0.89). We did not find any significant correlation between the degree of directional constraint and AQ scores in the separate analysis for each group.

**Figure 6.**
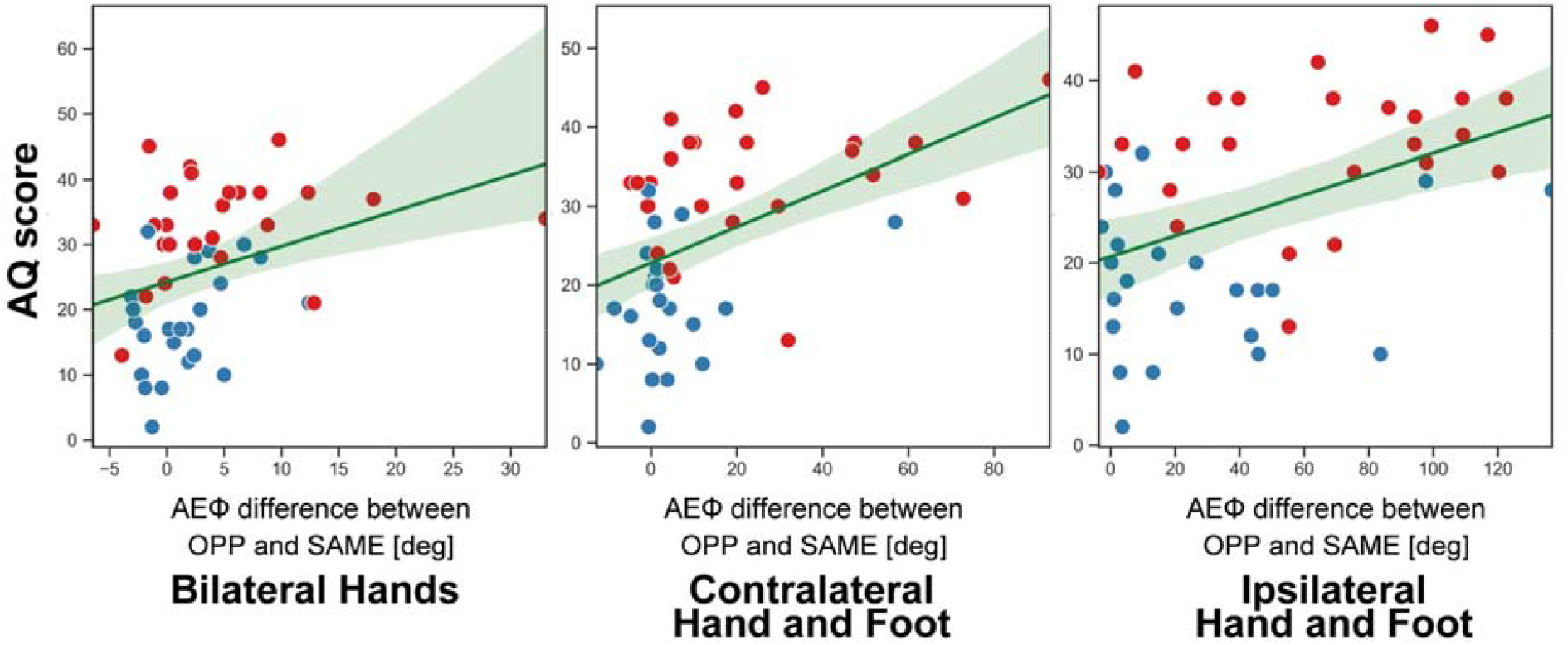
Correlation between the degree of directional constraint and AQ scores. The shaded areas (green, red, and blue for ASD + TD, ASD, and TD, respectively) denote the 95% confidence intervals for the correlation. The horizontal axis indicates the degree of directional constraint in each of the three limb combinations (bilateral hand, contralateral hand and foot, and ipsilateral hand and foot). The vertical axis indicates the AQ scores. Abbreviations: ASD, autism spectrum disorder; TD, typically developing; AQ, Autism Spectrum Quotient; AEФ, absolute error of the relative phase; SAME, movement in the same direction; OPP, movement in opposite directions.

## Discussion

This study focused on the directional constraint of inter-limb movements as a candidate mechanism of the impairment in coordination movements observed in individuals with ASD. We compared the degree of directional constraint during the cyclic movement of two limbs between ASD and TD participants, and we confirmed that a larger degree of directional constraint was associated with lower motor skills. Specifically, kinematic data revealed that ASD participants showed an increased directional constraint in the contralateral and ipsilateral hand–foot combinations compared with TD participants. Moreover, we found that participants with a larger degree of directional constraint exhibited poorer motor performance, especially in the body coordination category of the BOT-2, which requires hand–foot coordination skills. We also found that participants who exhibited stronger directional constraints tended to show severe autistic traits, as assessed by AQ scores.

### A larger degree of directional constraint was observed in individuals with ASD

Our main finding is that individuals with ASD showed a larger degree of directional constraint, especially in the contralateral and ipsilateral hand–foot combinations. In individuals with ASD, the relative phase error increased only with movements performed in OPP directions, although there was no difference between the two groups for movements in the SAME direction. These results indicate that the degree of error in individuals with ASD increases based on movement direction, and that this increase is greater than that observed in TD subjects.

Previous studies have indicated that the PMA and SMA are essential regions involved in the mechanism underlying directional constraint. Specifically, the PMA has been found to be critically involved in the temporal processing of cyclic motor tasks, as indicated by the fact that suppression of the left PMA using burst transcranial magnetic stimulation (cTBS) decreased the timing accuracy during synchronized finger tapping with the right and left hand (Bijsterbosch, Lee, Dyson-Sutton, Barker, & Woodruff, 2011). In addition, Byblow et al. (2007) utilized a dual-coil paired-pulse TMS protocol and found that networks involving the dorsal premotor cortex (PMd)–M1 facilitate movements of the hand and foot in the SAME direction (Byblow et al., 2007).

Moreover, the SMA has been suggested to play an important role in bimanual tasks, which involve tight temporal coordination between the different actions performed by the hands (Obhi, Haggard, Taylor, & Pascual-Leone, 2002). In addition, the SMA was found to contribute to the inhibition of unintentional movements during voluntary motor control (Sumner et al., 2007; Wardak, 2011), which extends beyond the fundamental role of the SMA in the coordination of multiple limbs (Debaere et al., 2001). In terms of cyclic hand– foot movements, suppressing neural activity in the SMA region using repeated TMS increased the phase error of movements in OPP directions (Steyvers et al., 2003). By contrast, facilitating the SMA using anodal tDCS improved the accuracy and stability of anti-phase bimanual supination-pronation movements (Carter et al., 2015).

Based on these findings, it has been hypothesized that the PMA and SMA have opposite roles in the performance of inter-limb movements; in other words, the PMA may facilitate in-phase movements, while the SMA may inhibit the neural signals that lead to limb movements in the synchronized phase and contribute to the execution of anti-phase actions. People with ASD have been found to have dysfunctional neural activation of inhibitory systems, as suggested by abnormal GABAergic signaling (Pizzarelli & Cherubini, 2011). Kana et al. (2006) demonstrated that the inhibitory circuitry of people with ASD was atypically activated and less synchronized during tasks that required response inhibition depending on visual signals (Kana, Keller, Minshew, & Just, 2007). In addition, several studies have shown that GABA is present in lower concentrations in the brain of people with ASD (Gaetz et al., 2014; Puts et al., 2017). Moreover, our previous work suggested that a decreased GABA concentration in the SMA is related to lower motor coordination skills in individuals with ASD (Umesawa et al., 2020). Perkins et al. (2015) observed that, during action observation (i.e., hand gestures such as waving, pointing, and grasping), neural activation of the premotor cortex is increased in high-functioning individuals with autism (Perkins, Bittar, McGillivray, Cox, & Stokes, 2015). In addition, imbalanced resting-state functional connectivity was observed in the PMA, and this abnormal connectivity was associated with stereotyped behavior in individuals with ASD (Huang et al., 2018). According to these previous reports, it could be hypothesized that a poorer inhibitory signal from the SMA or an increased synchronized signal from the PMA causes a strong directional constraint in ASD participants.

In the contralateral hand–foot combination, individuals with ASD showed increased directional constraint compared to TD participants, who displayed the same degree of constraint even in the bilateral hand combination. Previous studies have suggested that the neural modulation that facilitates movements in the SAME direction would work strongly for ipsilateral limb combinations but would be less effective for contralateral limb movements (Baldissera et al., 2002; Byblow et al., 2007). A comparison of the brain activity elicited by hand–foot movements in the SAME and OPP directions showed that the SMA becomes more activated for ipsilateral than for contralateral actions (Nakagawa, Kawashima, Mizuguchi, & Kanosue, 2016). Nakagawa et al. (2016) hypothesized that this might reflect a difference in the resources needed to inhibit the neural networks that are involved in disrupting ipsilateral and contralateral hand–foot movements in the OPP directions (Nakagawa et al., 2016). Thus, it can be speculated that synchronized signals would be weakened in the contralateral hand–foot combination compared with the ipsilateral hand–foot combination. A previous meta-analysis revealed a reduction in the size of the corpus callosum in autism (Frazier & Hardan, 2009), thus suggesting the possibility that, in people with ASD, neural signals are not weakened even in the contralateral hand– foot combination due to aberrant connectivity between the left and right cerebral hemispheres.

### The degree of directional constraint was associated with body coordination skills

Participants who showed an increased degree of directional constraint manifested lower body coordination scores. We suggest that directional constraint disrupts the independent movement of the hand and foot, thus resulting in poorer body coordination skills in daily life. In addition, we observed a significant relationship between the BOT-2 scores for strength and agility and the degree of directional constraint. This is because the strength and agility category also comprises tasks that require the separate movement of limbs, such as push-ups, sit-ups, and side steps, which partly require coordinated movements. Moreover, the BOT-2 scores for manual coordination significantly correlated with the degree of directional constraint in the contralateral and ipsilateral hand–foot combinations, although this category does not include tasks that use the lower limbs. Difficulties in the bilateral coordination of hands and arms may be caused by the same factors that lead to difficulties in coordinating the upper and lower limbs. We did not observe any significant relationship between the scores for fine motor control and the degree of directional constraint. This is unsurprising given that the tasks in this category include the fine control of fingers and the coordination of visual and hand movements; hence, there is little or no overlapping with the abilities required for performing inter-limb coordinated movements. In the group analysis, the degree of directional constraint significantly correlated with the scores for manual coordination and body coordination only in the ASD group. As mentioned above, ASD participants showed a remarkable increase in directional constraint in the contralateral hand–foot combination, whereas TD participants showed a lower degree of directional constraint during ipsilateral hand–foot movements. In healthy individuals, the neural signals that synchronize hand–foot movements are weakened in contralateral movements (Baldissera et al., 2002; Byblow et al., 2007), whereas the signals may maintain a certain degree of strength in individuals with ASD. Therefore, correlations were observed only in the ASD group due to the deficits of neural inhibitory function observed in this population (Kana et al., 2007; Pizzarelli & Cherubini, 2011).

### Individuals with severe autistic traits exhibited stronger directional constraint

Significant correlations were observed between autistic traits and the degree of directional constraint in all limb combinations. It has been reported that the PMA involves a mirror neuron system, which has been suggested to function as a basic mechanism for social cognition (Rizzolatti, Fogassi, & Gallese, 2001). A previous study demonstrated that abnormal event-related desynchronization in the low beta band (12–20 Hz), which has been suggested to be an index of mirror neuron activity, is observed in the PMA and SMA individuals with severe autistic traits during action observation (Puzzo, Cooper, Vetter, & Russo, 2010). According to these previous reports, it is possible that there is a common neural base that contributes to the directional constraint of inter-limb movements and social cognition.

## Conclusion

Our present findings provide evidence that (1) individuals with ASD are more likely than healthy individuals to display synchronized movement direction during hand–foot cyclic activities (i.e., directional constraint); (2) a stronger degree of directional constraint during hand–foot movements is associated with poorer body coordination skills; and (3) individuals with severe autistic traits tend to show stronger directional constraint during two-limb coordination. These results suggest that people with autism may have difficulties in inhibiting the neural signals that synchronize the direction of inter-limb movements, resulting in motor disabilities of bodily coordination.

## Acknowledgements

We would like to thank I. Wang for their technical help. We also appreciate the help of A. Nakajima in participant recruitment.

## Ethics approval and consent to participate

All participants and their parents provided written informed consent after the study procedures were fully explained. The study was approved by the Ethics Committee of the National Rehabilitation Center for Persons with Disabilities.

